# Small Structural Variations, Large Functional Consequences: Comparative Analysis Reveals Structural Control of Ubiquitylation Site Selection by BRCA1/BARD1

**DOI:** 10.64898/2026.06.26.734868

**Authors:** Sahar Heidari, Caitlin Lightle, Lauren Herrington, Tejas Shah, Fariha Hossain, Faruck Morcos, Susan T. Weintraub, Mikaela D. Stewart, Hedieh Torabifard

## Abstract

BRCA1/BARD1 is a chromatin-associated E3 ubiquitin ligase that ubiquitylates histone H2A to coordinate DNA damage repair, transcriptional repression, and genome stability. In *Caenorhabditis elegans (C. elegans)*, the orthologous BRC-1/BRD-1 complex performs analogous functions but exhibits structural variation, most notably through an additional 11-residue loop in BRD-1 that is absent from human BARD1. Prior experiments indicate this worm-specific insertion promotes nucleosome engagement and may alter the preferred lysine target for ubiquitylation. Here, we provide a cross-species comparison by integrating computational and experimental investigation to clarify how a discrete structural variation can tune BRCA1-family ligase behavior and, consequently, chromatin regulation. *In vitro* ubiquitylation assays and mass spectrometry reveal BRC-1/BRD-1 ubiquitylate the C-terminal tail of histone H2A with less specificity than the human homologs. All-atom molecular dynamics simulations of both the *C. elegans* BRC-1/BRD-1–LET-70-Ubiquitin assembly and the human BRCA1/BARD1–UbcH5c-Ubiquitin complex in the presence of the nucleosome core particle uncover that the BRD-1 loop makes transient contacts with nucleosomal DNA and histone tails, thereby modulating the positioning and conformational flexibility of the bound E2 (ubiquitin-conjugating enzyme). Together, our results suggest that the BRD-1 loop alters the E3–E2 geometry, thereby altering ubiquitylation-site specificity.

## Introduction

Breast and ovarian cancer risk is strongly influenced by inherited mutations in the tumor-suppressor gene, BRCA1 and its partner, BARD1.^1,2^ The N-terminal RING domain of BRCA1 pairs with the RING domain of BARD1 to form the BRCA1/BARD1 (BCBD) heterodimer, an E3 ubiquitin ligase^3–5^ central to genome maintenance across diverse tissues.^6,7^ Through this partnership, BRCA1 contributes to DNA repair, cell-cycle checkpoint control, and transcriptional regulation. ^3^ Given that failures in genome maintenance elevate cancer susceptibility, understanding the underlying E3 mechanism and substrate control has received intense attention.^8–10^

A key BCBD activity is ubiquitylation of nucleosomal histone H2A at defined C-terminal lysines, a modification associated with transcriptional repression and DNA damage responses.^3,9,11–13^ While the BRCA1 RING domain contains a majority of the E2 (ubiquitin-conjugating enzyme) binding interface, BARD1 is not merely an accessory to BRCA1. The BARD1 RING has two established functions required for efficient ubiquitination of nucleosomal substrates. First, a positively charged residue at BARD1 position 99 interacts with ubiquitin (Ub) to position it for transfer. ^9,14^ Second, similar to BRCA1, specific residues on BARD1 contribute to nucleosome engagement. In human BCBD, BRCA1 contacts the H2A/H2B acidic patch through Lys70 and Arg71, whereas BARD1 engages the H2B/H4 cleft via Pro89 and Trp91.^13,14^ Although the structure of nucleosome-bound BCBD shed light on these critical contacts, ubiquitin and disordered histone tails are not present in the structure, leaving the precise functional contributions of BARD1 to lysine site selection and catalytic efficiency enigmatic. ^12^

Addressing these unknowns *in vivo* is challenging in mammals, as complete BRCA1 or BARD1 deficiency prevents viable embryonic development. By contrast, *Caenorhabditis elegans* (*C. elegans*) provides a powerful comparative model to probe the open questions *in vivo*. *C. elegans* is a free-living, non-parasitic, self-fertilizing nematode, often called “the worm”, that has long been a model for studying animal development and reproduction, with many pathways conserved across eukaryotes. ^15,16^ It provides a tractable *in vivo* alternative to human tissue studies because it is easy to cultivate, develops rapidly, remains transparent throughout its life cycle, and is highly amenable to genetics. ^17–19^ *C. elegans* shares genetic conservation with humans, including orthologs of *BRCA1* and *BARD1*, called *brc-1* and *brd-1*. The domain architecture for the two proteins encoded is similar across species; both have N-terminal RING domains with sequence similarity of 59% for BRCA1/BRC-1 and 72% for BARD1/BRD-1. ^14^ Consistent with this, the BRC-1/BRD-1 complex is found on meiotic chromosomes, participates in DNA damage responses, and ubiquitylates chromatin-associated substrates after DNA damage.^14,20–22^

These findings support conserved contacts and functional similarities between human and worm complexes. Although prior work^14,23^ implicates BRD-1 in ubiquitylation like its human counterpart BARD1, its precise mechanistic role and contribution to site specificity remains unclear. A key open question is the specificity of H2A ubiquitylation. In humans, BCBD preferentially modifies C-terminal lysines on disordered nucleosomal H2A. ^11^ For *C. elegans*, the targeted lysine(s) and the determinants of site selectivity have not been fully established, leaving it unclear whether *C. elegans* ubiquitylation mirrors human specificity. In this study, we integrate experimental and molecular dynamics (MD) approaches to identify the modified C-terminal lysines on *C. elegans* H2A and to define the determinants governing site-specific H2A ubiquitylation by the BCBD complex. Comparative analysis of human BCBD (HsBCBD) and *C. elegans* BCBD (CeBCBD) in the presence of the nucleosome core particle (NCP) highlights the role of BARD1/BRD-1 in substrate targeting.

## Methods

### Generating H2A lysine mutant constructs

Potential lysine targets on histone H2A were mutated to residues that cannot be ubiquitylated to determine which lysine(s) is targeted by the CeBCBD. K119 and K120 mutations in the H2A C-terminal tail were obtained using the QuikChange Site-directed Mutagenesis protocol with the modifications as described previously.^24^ The template for mutagenesis contained hybrid H2A with the sequence of *C. elegans* C-terminus substituted on the human H2A core in a pHis vector and already containing the K125H mutation tested and published previously.^14^ Separate PCR reactions were run containing either the single forward (ATCCAGGCGGTTCTGCTGCGTCGTACCGGAGGAGAC) or reverse primer (complement to the forward) to substitute both positions simultaneously using an annealing temperature of 58°C. The products were then melted and annealed together before a two-hour Dpn1 digestion at 37°C to cleave the parental template. N-terminal mutations of K13 and K15 were obtained using the New England Biolabs (NEB) Q5 Site-directed mutagenesis protocol using both unmutated hybrid H2A and hybrid H2A lacking C-terminal lysine residues as templates. Substitution of both lysine residues was achieved with an annealing temperature of 68°C and a single set of forward and reverse primers designed using the NEBasechanger online tool (GCGTTCTCGTTCTTCTCGTGCGG and GCACGCGCACGCGCTTTACCACC, respectively). Genetic sequencing using primers complementary to the T7 terminator or T7 promotor sequence was used to confirm the presence of the mutations.

### Nucleosome core particle reconstitution and protein purification

Specific ubiquitylation of target H2A lysines is only achieved when assays are carried out in the context of the nucleosome. ^11^ In order to generate nucleosomes containing the H2A chimera mutants, plasmids were sent to Sam Witus in Rachel Klevit’s laboratory at the University of Washington, where the histones were overexpressed in *E. coli* and used to construct histone octamers with human H2B, H3, and H4 as delineated previously.^12^ Octomers were shipped to Texas Christian University overnight on dry ice and stored at -80°C until needed for assays. Reconstitution of the octomers into nucleosomes was achieved through slow dialysis to remove salt in the presence of DNA as described previously.^12^ Briefly, 7 *µ*M of Widom 601 DNA and 8 *µ*M of the octamers were dialyzed in a solution containing 2 M

NaCl and 20 mM TRIS. The NaCl solution was diluted with 20 mM TRIS to a final salt concentration of 180 mM over the course of 36 hours to allow the octamers to bind DNA. Nucleosome core particles (NCPs) formed at a salt concentration of approximately 400 mM NaCl and were then subjected to a second dialysis in 20 mM TRIS and 50 mM NaCl for storage. Reconstitution was confirmed through TBE gel electrophoresis. Nucleosomes were stored on ice at 4 °C. Ub, LET-70 (E2), UBA1 (E1) enzyme, and the RING domains of BRC-1 and BRD-1 were individually overexpressed from a DNA plasmid in BL21(DE3) *E. coli* cells and then the proteins were isolated via column chromatography. Specific construct links used as well as purification techniques were published previously.^14^

### Nucleosome assays

Nucleosome assays were used to measure CeBCBD E3 ligase activity in the presence of lysine mutations on the H2A chimera. The reactions each contained 25 mM HEPES, 150 mM NaCl, 0.3 *µ*M nucleosomes, 20 *µ*M Ub, 0.5 *µ*M human UBA1 (E1), 4 *µ*M LET-70 (E2), 5 mM ATP, 5 mM MgCl_2_, and 8 *µ*M CeBCBD (E3). The nucleosomes used were either wild-type or contained one or more lysine mutations in the H2A chimera. The assays were run at 37 °C and 400 rpm shaking, with samples taken at 0, 10, and 30 minutes and added to 2x load dye containing sodium dodecyl sulfate (SDS) to denature the proteins and stop the reaction. The zero time point was taken prior to the addition of ATP. Time points taken from the nucleosome assays were analyzed via western blotting and then quantified using ImageJ.^25^ The samples were first run on a 15% polyacrylamide gel and then transferred to a nitrocellulose membrane. The samples were blocked with non-fat dry milk proteins at a 5% concentration and blotted using a 1:5000 dilution of a rabbit primary antibody to detect the VSVG-tag on histone H2A (Millipore Sigma Corporation) followed by a 1:10000 dilution of a goat anti-rabbit secondary antibody (Rockland Immunochemicals Incorporated). The H2A bands were visualized via a BCIP/NBT kit (Promega Corporation) according to the manufacturer’s instructions. The western blots were quantified using ImageJ, comparing unmodified H2A within the sample at each time point to the zero time point.^25^ Means from two replicates of the assay were compared to wild-type using a Student’s t-test, and lack of statistical significance was determined at both 10 and 30 min given p-values *>* 0.1 for all mutants.

### Mass spectrometry

In preparation for mass spectrometry, a large-scale (72 *µ*L) nucleosome assay was performed using the CeBCBD E3 ligase on wild-type H2A chimera. This was done using the same procedure as the nucleosome assays described previously with a few modifications of increased protein concentrations (nucleosomes at 3.7 *µ*M, CeBCBD at 24 *µ*M, LET-70 at 7 *µ*M, E1 at 0.6 *µ*M, Ub at 71 *µ*M, and ATP at 5.9 *µ*M) and the assay was allowed to run for one hour to maximize ubiquitylation of H2A. An aliquot of the sample (40 *µ*L) was mixed with 10 *µ*L 12% phosphoric acid and 50 *µ*L 10% SDS/50 mM triethylammonium bicarbonate (TEAB), applied to S-Traps (mini; Protifi), reduced/alkylated with a mixture of tris(2-carboxyethyl)phosphine hydrochloride (TCEP) and 2-chloroacetamide and digested for 2 hr at 30 °C with chymotrypsin (sequencing grade; Sigma) in 50 mM triethylammonium bicarbonate (TEAB). Peptides were eluted from the S-Traps sequentially with 50 mM TEAB, 0.2% formic acid, and 0.2% formic acid in 50% aqueous acetonitrile. The pooled eluates were dried by vacuum centrifugation and redissolved in 30 *µ*L of the starting mobile phase (see below). Digests were analyzed by capillary HPLC-electrospray ionization tandem mass spectrometry on a Thermo Scientific Orbitrap Fusion Lumos mass spectrometer. On-line HPLC separation was accomplished with an RSLC NANO HPLC system (Thermo Scientific/Dionex) interfaced with a Nanospray Flex ion source (Thermo Scientific) using the following specifics: column was a PepSep (Bruker; ReproSil C18, 15 cm x 150 *µ*m, 1.9 *µ*m beads); mobile phase A was 0.5% acetic acid (HAc)/0.005% trifluoroacetic acid (TFA) in water; mobile phase B was 90% acetonitrile/0.5% HAc/0.005% TFA/9.5% water; gradient was 3% to 42% B in 60 min; flow rate was 800 nL/min. Precursor ions were acquired in the Orbitrap in centroid mode at 120,000 resolution (*m/z* 200); data-dependent higher-energy collisional dissociation (HCD) spectra were acquired at the same time in the linear trap using the “rapid" speed option and 30% normalized collision energy. Other MS scan parameters included: mass window for precursor ion selection, 0.7; charge states, 2 – 5; dynamic exclusion, 15 sec (± 10 ppm); intensity to trigger MS2, 50,000. For peptide analysis, Mascot (v2.8.3; Matrix Science, London UK) was used to search the spectra against a combination of the following databases: UniProt_Human_reference [UniProt_Human_ref 9606_20220216] (20,588 sequences; 11,394,277 residues); an in-house database that includes the sequences of recombinant and target proteins (773 sequences; 329,786 residues); and common contaminants (not including *Bos taurus* proteins, (124 sequences; 62,564 residues)). Cysteine carbamidomethylation was set as a fixed modification while Arg-Gly-Gly-Lys, methionine oxidation, and deamidation of glutamine and asparagine were considered as variable modifications; chymotrypsin was specified as the proteolytic enzyme, with three missed cleavages allowed. The Mascot search results were imported into Scaffold (version 5.3.3, Proteome Software Inc., Portland, OR). Acceptance of peptide and protein identifications were based on the following: peptides with greater than 90.0% probability by the *Percolator* posterior error probability calculation^26^ and proteins with greater than 99.0% probability based on the *Protein Prophet* algorithm.^27^ A minimum of two identified peptides was required. These settings resulted in a protein-level false discovery rate of *<* 0.01%. Proteins that contained shared peptides and could not be differentiated based on tandem-MS analysis alone were grouped to satisfy the principles of parsimony.

### Molecular dynamics simulations

#### Model preparation

The cryo-EM structure of human BRCA1/BARD1–UbcH5c bound to the NCP (PDB ID: 7JZV^12^) was used to generate the starting structure. Missing histone-tail residues were built using SWISS-MODELLER,^28^ and Ub was subsequently attached. A Ub monomer was isolated from human tetraubiquitin (PDB 1F9J^29^) in VMD,^30^ and was positioned with its C-terminal Gly76 near the E2 active site. An isopeptide bond between Ub’s C-terminal Gly76 and the *ε*-amine group of the active-site lysine (E2-K197) was created using tleap module from the AmberTools20;^31^ all other system components matched the previously published Ub-free model.^32^ Owing to the E2 active-site substitution in PDB,^12^ this model features an E2∼Ub isopeptide bond rather than the native thioester intermediate.^33,34^

For *C. elegans*, we used the same 7JZV coordinates from the human complex as a template. The BRC-1 RING domain, BRD-1 RING domain, and LET-70 were modeled simultaneously using 7JZV as a homology model using the SWISS-MODEL web sever. ^28^ The *C. elegans* H2A (UniProt P09588) was similarly modeled on the human template using the SWISS-MODEL web server, and the three missing H2A C-terminal tail residues were added using PyMOL.^35^ Ub sequence alignments indicated a single proline (PRO19) to alanine substitution in worm relative to human; accordingly, we mutated that residue while retaining the same Ub coordinates. Protonation states of all components at pH 7.4 were assigned with H++.^36^ We also protonated the BRD-1 histidine corresponding to BARD1 R99 to preserve a comparable positive charge.^14^ This model is shown in Figure 1a.

**Figure 1:**
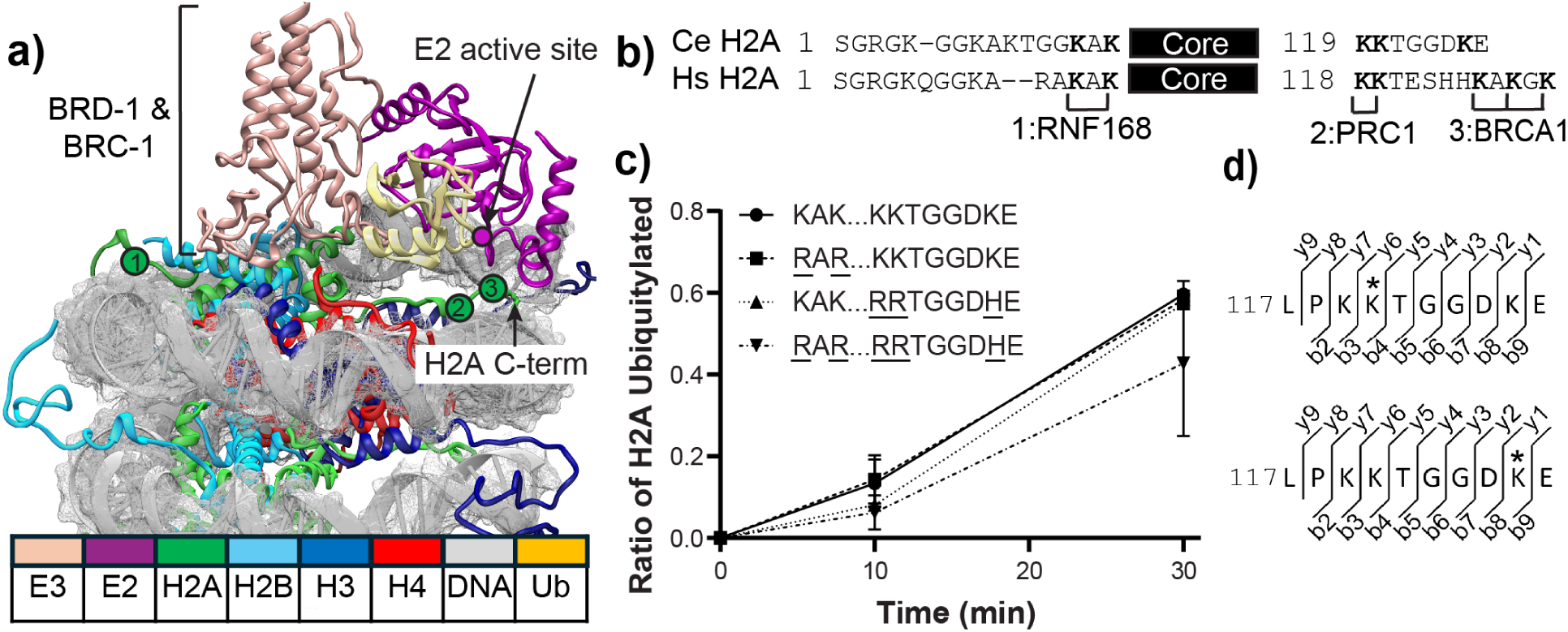
CeBCBD is not specific for the most C-terminal Lys on H2A. a) Model of *C. elegans* nucleosome ubiquitylation complex built from the cryo-EM structure of the human complex 7JZV. Areas containing conserved H2A lysine residues (highlighted with green circles labeled 1-3) are not in close enough proximity to the E2 active site (magenta circle) for ubiquitylation. b) Alignment of the N- and C-terminal tail sequences of H2A from *C. elegans* (Ce) and *H. sapiens* (Hs) to show the sites of ubiquitylation established for human RNF168, PRC1, and BRCA1 are semi-conserved in worms. Numbering (1-3) refers to the conserved area of the H2A structure highlighted with green circles in a. c) Mutation of semi-conserved lysine residues does not significantly diminish H2A ubiquitylation by CeBCBD *in vitro*. The first three amino acids represent the conserved RNF168 site on the H2A N-terminus, while the remaining amino acids represent the H2A C-terminus, with the core represented by ellipses. d) Mass spectrometry detects ubiquitylation of the C- terminal tail of H2A (residues 117-126) by CeBCBD. Hatched marks above and below the peptide sequence correspond to the detected b and y fragment ions, respectively, in the MS/MS. Asterisks mark the location of Ub addition as noted by the extra mass corresponding to the C-terminus of Ub residues after chymotrypsin cleavage (RGG).

All systems were solvated with TIP3P water, ^37^ neutralized with Na^+^, and adjusted to a physiological salt concentration of 150 mM by adding NaCl in tleap. ^31^ The force fields ff14SB,^38^ DNAOL15^39^ and ZAFF^40^ in AMBER were used to model the protein, nucleic acids and tetrahedrally coordinated zinc atoms, respectively. The final human and *C. elegans* systems contained ∼406,000 and ∼378,000 atoms, respectively.

#### Simulation details and analysis

Systems were relaxed with a two-stage minimization: first, 3,000 steps of steepest descent followed by 2,500 steps of conjugate gradient with harmonic restraints (500 kcal·mol*^−^*^1^·Å*^−^*^1^) on the protein and nucleosome; and second without restraints. After minimization, systems were heated in three stages over 1.5 ns from 0 to 300 K under NVT with a Langevin thermostat.^41^ The systems were then equilibrated for 1 ns under NPT with weak backbone restraints (10 kcal·mol*^−^*^1^·Å*^−^*^1^) and 100 ns afterwards without any restraints. Production simulations were run in triplicate for 800 ns each in the NPT ensemble with frames saved every 50 ps to ensure adequate sampling. All NPT simulations were performed by controlling the temperature at 303 K using Langevin thermostat^41^ and pressure at 1 atm using Berendsen barostat.^42^ All simulations were carried out using the GPU-accelerated pmemd.cuda engine in AMBER20 package.^43^ SHAKE algorithm^44^ was applied to constrain bonds involving hydrogen atoms. Long-range electrostatics were treated with particle-mesh Ewald, ^45^ and a 12 Å cutoff was used for non-bonded interactions. Simulation input and parameter files for both systems are available at Zenodo.

Root mean square displacement (RMSD) of backbone atoms and root mean square fluctuation (RMSF) of side chains were calculated for each trial with reference to the first frame and the average structure, respectively, to assess structural dynamics. The average RMSD results in Figure S1 shows that all the components are equilibrated in 100 ns. Therefore, all analyses were performed using AmberTools20’s cpptraj module^46^ and VMD^30^ on 800 ns of production trajectories after the system had equilibrated for 100 ns. The B-factor of BARD1 and BRD-1 was derived from the RMSF results. We combined all replicate production trajectories and performed contact-frequency analysis, tilt-angle measurements, and principal component analysis (PCA). Contact-frequencies were computed in VMD using an in-house Tcl script and a 4.5 Å distance cutoff between residues. For E3 tilt-angles, we defined the angle between a vector along the H2B *α*-helix and a vector along the *α*-helix of BRCA1/BRC-1 or BARD1/BRD-1, yielding the BRCA1/BRC-1 and BARD1/BRD-1 tilt angles, respectively. PCA was conducted to cluster the motions, and the resulting projections were partitioned into discrete states for the human and *C. elegans* systems. Distance analyses were performed for each *C. elegans* state and across all human replicates to measure the distance between the E2 active site and the *ε*-amine group of the target lysine residue(s) on the H2A tail. Visualization of the simulations as well as figure generation, was performed using VMD,^30^ PyMOL^35^ and ChimeraX^47^ packages.

### Coevolutionary Paralog Pairing and Comparative Length Analysis Across Nematodes and Mammalia

HMMER software uses profile Hidden Markov Models (HMMs) to identify homologous sequences across diverse organisms.^48^ From a single sequence or a small initial multiple sequence alignment (MSA) of related family members, the *HMMSearch* tool generates a larger, more comprehensive MSA from a profile HMM.

The BARD1/BRD-1 domain occurs in many systems, but the combination of BRCA1-BARD1 pairing is distinctive and allows for more targeted homology searches. An initial MSA of the full BRCA1-BARD1 complex from *H. sapiens* (Q99728) and *C. elegans* (Q21209) was used as seed to build a broader alignment via HMMER. This alignment was refined by removing highly gapped sequences (consecutive gapped regions *>* 15% of sequence length) and ensuring that RING domain in BARD1/BRD-1 region retained the canonical *CX*_2_*CX*_(9_*_−_*_39)_*CX*_(1_*_−_*_3)_*HX*_(2_*_−_*_3)_*CX*_2_*CX*_(4_*_−_*_48)_*CX*_2_*C* (X is any amino acid) motif along with a positively charged residue (R,K,H) at MSA aligned equivalent of position 99. ^9,14,49^ Similarly, an initial MSA of H2A from *H. sapiens* (P20671) and *C. elegans* (P09588) was used as seed to create a larger alignment using HMMER. This was further refined by removing highly gapped sequences (consecutive gapped regions *>* 15% of sequence length) and removing sequences that did not contain the H2A signature [AC]-G-L-X-F-P-V (X is any amino acid) pattern (PROSITE entry PS00046).^50^ Finally, sequences were taxonomically classified using the organism identifier (OX code) embedded in each FASTA header, parsed with the ETE toolkit.^51^ Only sequences belonging to Nematoda/Chromadorea (worms) and Mammalia (mammals) were retained for downstream analysis.

Although each species encodes multiple H2A isoforms, ubiquitylation by BARD1 has shown to target lysine (K) residue(s) in the C-terminal tail, corresponding to the last 10 residues following the conserved double lysine in human H2A (K118 and K119; Figure S2).

Thus, H2A isoforms lacking additional lysine residues in this C-terminal region and equivalent positions defined by the alignment in other species were excluded from further analysis. Even after these refinements, both the BRCA1-BARD1 and H2A alignments retained multiple paralogous copies for many species. Biologically relevant interacting pairs among these paralogs were identified through coevolutionary analysis. For each species with two interacting proteins, residues are evolutionary coupled driving their physical interaction. Direct Coupling Analysis (DCA) exploits this signal by statistically quantifying direct coupling between pairwise residue positions across two proteins, allowing us to predict biologically meaningful interacting partners.^52^ The paralog pair matching algorithm developed by Gueudré *et al*., which utilizes two MSAs as input and uses DCA-derived inter-protein co-evolutionary signal scores to infer the optimal pairing between sequences. ^53^ For each species in which both interacting proteins are present, we utilized this algorithm to identify the combination of paralog pairs that maximizes the inter-protein coupling score, producing a concatenated MSA that reflects the most likely biologically interacting partners.

Using the concatenated MSA, we examined two structural features for each species: the number of non-gap residues in the H2A C-terminal tail (corresponding to the last 10 residues of human H2A) and the length of the BRD-1 loop insertion relative to *C. elegans*, both mapped to alignment equivalent positions. The relationship between these features was visualized using a jittered scatter plot, where small random offsets are added to reduce overlapping data points, revealing strong reciprocal correlation between worms (Nematoda/Chromadorea) and mammals (Mammalia).

## Results

### Target lysine specificity is not conserved between CeBCBD and Hs-BCBD

Previous mutagenesis studies suggested the lysine on H2A targeted by the human and worm BCBD heterodimers may vary.^14^ Mutation to remove the three most C-terminal lysine residues of H2A severely limits the activity of the HsBCBD *in vitro* (labeled “BRCA1" in Figure 1b).^11,54^ However, mutation of the Ce H2A lysine closest to the positions targeted by HsBCBD did not significantly alter ubiquitylation of CeBCBD. ^14^ To determine the target of *C. elegans* BRC-1/BRD-1 complex, we first modeled the CeBCBD structure bound to the nucleosome using the cryo-EM structure of HsBCBD (7JZV) as a homology model (Figure 1a). Minimization of this structure does not result in H2A lysine residues in close enough proximity to the E2 active site to gauge which will be modified.

In further attempts to locate the H2A ubiquitylation site, we mutated all conserved lysine residues on the H2A tails and conducted *in vitro* ubiquitylation assays. While the folded core of histone H2A has 98% sequence similarity between humans and worms, the tail sequences are more divergent (Figure 1b, core sequences shown in Figure S2). To identify the alternate or additional site(s) of ubiquitylation, we incorporated mutant H2A into reconstituted nucleosomes *in vitro* and assayed them using western blotting for an N-terminal epitope tag on H2A. The shift in molecular weight upon ubiquitylation results in distinct migration patterns of the unmodified and ubiquitylated H2A (Figure S3), which allows for quantification of the total amount ubiquitylated for each mutant (Figure 1c). These assays reveal CeBCBD ubiquitylation of H2A was not significantly impacted by elimination of the lysine residues targeted by human RNF168 (squares), any of the C-terminal lysine residues (triangles), nor all of these residues mutated simultaneously (inverted triangle).

When the mutation site could not be determined via rational mutagenesis, we digested the modified H2A in the absence of lysine mutations and used tandem mass spectrometry (MS/MS) to determine the location of ubiquitylation. We found evidence of ubiquitylation at two sites on the C-terminal H2A tail (Figure 1d and S4). While similar experiments using HsBCBD produced only modification of the far C-terminal lysine residues K127 and K129, ^11^ here we show CeBCBD is not specific for the far C-terminal lysine residue(s) given K120 ubiquitylation is also detected as depicted in Figure 1d.

### BARD1 and BRD-1 interact with the nucleosome through non-conserved interfaces

To understand why CeBCBD is less specific for far C-terminal Lys compared to HsBCBD, we examined their structural differences. As in humans, the *C. elegans* orthologs BRC-1 and BRD-1 contain conserved RING domains that confer E3 ubiquitin ligase activity. ^14^ BRC-1 is largely structurally conserved (lighter colored domain in Figure 2a) and its nucleosome binding site was also previously shown to be conserved.^14^ However, BRD-1 contains an additional 11 residues relative to BARD1, which may influence the architecture, interactions, and dynamics of the complex. As shown in Figure 2a, alignment of the minimized HsBCBD and CeBCBD structures highlights an extra loop corresponding to the 11 additional residues in BRD-1. To further investigate the role of this loop, we conducted MD simulations on the nucleosome-bound complexes and analyzed the RMSD, RMSF, RMSF-derived B-factors and tilt angle for BRD-1 and BARD1 from the simulations. We assessed the overall structural stability of each E3 by computing the backbone RMSD. Comparison of RMSD between the human and *C. elegans* E3s in Figure S5a shows that BRD-1 exhibits higher RMSD values and a broader distribution across replicates than BARD1, indicating greater conformational mobility in BRD-1, whereas BRC-1 and BRCA1 display RMSDs in a similar range. Per-residue RMSF profiles further support this pattern: BRD-1 shows larger fluctuations than BARD1 overall, with the greatest amplitudes in the BRD-1 extra-loop, shown in Figure S5b, while BRC-1 and BRCA1 shows similar fluctuations. The B-factor results in Figure 2b visually reflect this fluctuation by highlighting the pronounced mobility of the BRD-1 extra loop. The tilt angle analysis quantifies the orientation of the BCBD heterodimer relative to the nucleosome surface that contains the H2A C-terminal tail. Larger angles indicate a lean toward this surface; smaller angles indicate a tilt away. Tilt angle between BCBD and the nucleosome showed that CeBCBD samples a wider range of positions relative to the nucleosome than HsBCBD. As shown in Figure 2c, BRD-1 explores tilts of ∼20*^◦^* to ∼140*^◦^* (span ∼120*^◦^*) whereas BARD1 covers ∼30*^◦^* to ∼100*^◦^* (span ∼70*^◦^*). This broader coverage indicates larger movement for BRD-1, consistent with the BRC-1 vs BRCA1 tilt angle difference shown in Figure S6b; although BRC-1 tilts more than BRCA1 (likely due to its coupling to BRD-1), this difference is smaller than that between BRD-1 and BARD1. The wider tilt-angle span of BRD-1 is consistent with the RMSD and RMSF trends, reinforcing increased mobility of BRD-1 in the nucleosome-bound state.

**Figure 2:**
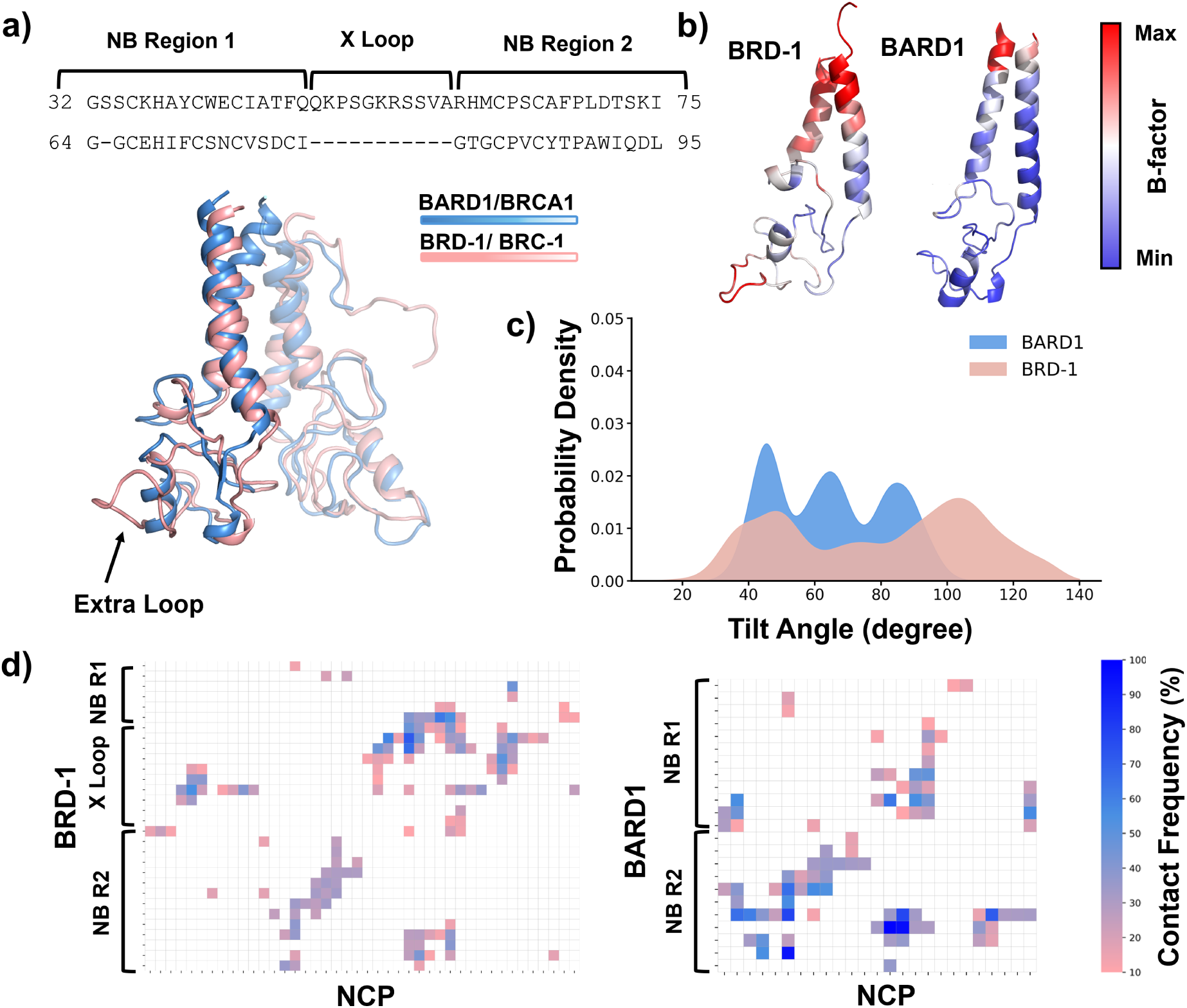
BARD1 and BRD-1 interact with the nucleosome using non-conserved binding interactions. a) Comparisons of minimized structures (nucleosome-bound HsBCBD (PDB ID: 7JZV^12^) vs minimized structure of nucleosome-bound CeBCBD) and sequence alignment of BARD1 and BRD-1 nucleosome binding (NB) regions categorized as “NB region 1”, “NB region 2” and extra-loop (“X Loop”). b) Differences in BARD1 and BRD-1 flexibility assessed by B-factor analysis derived from RMSF values and normalized to 1-100 scale. Red indicates the maximum B-factor (highest flexibility), and blue indicates the minimum B-factor (lowest flexibility). c) Tilt-angle analysis for BARD1 and BRD-1 relative to the nucleosome surface. d) Differences in BARD1 vs BRD-1 nucleosome contacts over the course of simulation, analyzed by contact frequency. For ease of comparison, residues contacting the nucleosome were categorized as nucleosome binding regions 1 and 2 (“NB R1” and “NB R2”) and the extra loop of BRD-1 (“X Loop”).

To further assess the contribution of BRD-1 extra loop in the dynamics of these complexes, contact-frequency patterns were analyzed by comparing the nucleosome-E3 ligase contacts in human and *C. elegans* complexes. CeBCBD forms a greater number of unique contacts with NCP in comparison to HsBCBD, although the human contacts persist through more frames. Figure S6c indicates that BRCA1 and BRC-1 exhibit similar contact profiles, with subtle deviations arising from differences in their dynamics and tilt extent. In contrast, Figure 2d shows that the major differences originate from BRD-1 and BARD1. BRD-1 engages more residues in nucleosome contacts that are more broadly distributed across the nucleosome but individually less persistent, whereas BARD1 forms fewer interactions, many of which remain stable throughout most of the simulation. Importantly, residues within the extra loop in BRD-1 display relatively persistent interactions with NCP as shown in Figure 2d. The specific BRD-1 and BARD1 residues that contact the NCP are delineated in Figure S7. Therefore, the localized flexibility in BRD-1 extra loop may contribute to species-specific differences in E3 ligase dynamics and, in turn, ubiquitylation. Together, these results support the hypothesis that the additional loop in BRD-1 contributes significantly to the different dynamic behavior of the *C. elegans* and human complexes.

### CeBCBD adopts more diverse nucleosome binding configurations, enabling targeting of additional lysines compared to HsBCBD

Differences in the intrinsic dynamics of BRD-1 and BARD1 are likely to be transmitted to the overall motion and behavior of the E3–E2 complex. Because the E2 enzyme is covalently linked to Ub and plays a central role in catalyzing Ub transfer, changes in the positioning and flexibility of BRD-1 versus BARD1 will alter the orientation and conformational ensemble of the E2∼Ub conjugate in the *C. elegans* and human complexes. These shifts in E2∼Ub geometry, which are coupled to E3 motion, can, in turn, modulate the efficiency and specificity of ubiquitylation. To gain additional insight into differences in the structural conformations sampled by the *C. elegans* and human complexes, PCA was performed on BCBD with respect to the NCP. PCA was conducted on all trials together. The first two principal components capture two distinct motions. *Tilt-1* and *Tilt-2* motions represent the first principal component (PC1) and second principal component (PC2), respectively. *Tilt-1* corresponds to tilting of the BCBD toward H2B, whereas *Tilt-2* corresponds to a motion that drives E2 towards the nucleosome surface containing H2A C-terminal tail. Figure S8 provides a schematic representation of the *Tilt-1* and *Tilt-2* motions.

Because ubiquitylation occurs when the E2∼Ub conjugate approaches the C-terminal tail of H2A, motion along *Tilt-2* (PC2) most directly tunes the catalytically competent geometry of the complex. Analysis of the PC2 reveals three distinct conformational states (Figure 3a) that differ in the extent to which the E2∼Ub module is oriented toward the H2A tail.

**Figure 3:**
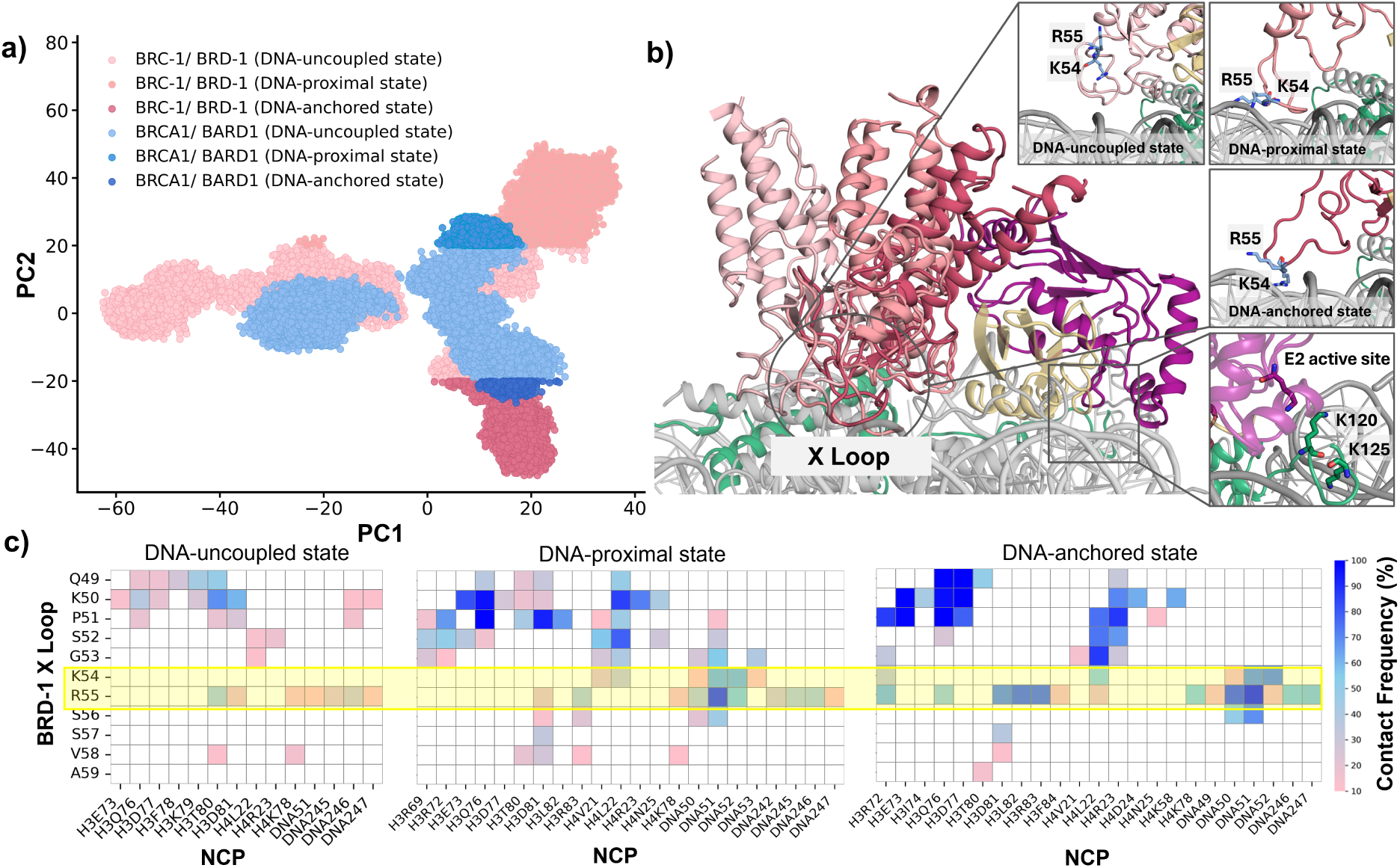
Principle component analysis reveals three predominant bound states, dictated in part by BRD-1 contacts with DNA. a) PCA of CeBCBD and HsBCBD. Pink indicates the *C. elegans* complex and blue the human complex. PC2 describes a motion that pushes E2 towards the nucleosome surface containing H2A C-terminal tail, revealing three states for the *C. elegans* and a single dominant state for the human complex. The PC2 cutoff used to define these states was selected based on visualization of the trajectories. b) Different orientations of CeBCBD and its extra loop corresponding to the three observed states. c) Contact frequencies between the BRD-1 extra loop and the NCP vary across the states.

These states, therefore, provide a structural framework for understanding how E3-driven tilting of the complex can regulate access of E2∼Ub to its lysine targets on H2A. While the overall movement patterns are similar for both complexes, CeBCBD explores a broader range of motions than HsBCBD, reflecting the greater flexibility of the *C. elegans* complex. The additional loop in BRD-1 plays a central role in this behavior, contributing to distinct conformational states that are absent or substantially attenuated in the human system. Specifically, CeBCBD samples all three states, termed DNA-uncoupled, DNA-proximal and DNA-anchored states, with pronounced motions, whereas HsBCBD predominantly occupies DNA-uncoupled state, with the other two states being much less evident.

Figure 3b presents a representative conformation of the *C. elegans* states and the factors driving these motions. Interactions between K54 and R55 in the BRD-1 loop and DNA drive worm-specific dynamics and the distinct motion patterns detected. Figure 3c shows frequent contacts between the BRD-1 loop and the NCP at different states, highlighting the importance of these interactions in shaping movements within the *C. elegans* complex: in DNA-uncoupled state, the interactions between K54 and R55 with DNA are absent/negligible, consistent with this state being the most prevalent in the human complex when the loop is absent; in DNA-proximal state, loop interactions are intermediate; and in DNA-anchored state, they are most frequently observed. Linear interaction energy analysis further supports this observation, as the sum of electrostatic and van der Waals contributions in Figure S9 indicate the increase in interactions of K54 and R55 with the nucleosome which are primarily mediated by DNA rather than by histone contacts. The interactions of these two positively charged residues increase the number of contacts between the extra loop and the NCP; accordingly, Figure 3c shows that Q49, K50, and P51 make progressively more stable contacts with the histones from DNA-uncoupled to DNA-anchored state.

In DNA-anchored state, where CeBCBD bends toward the H2A C-terminal tail, K54 and R55 form stable contacts with the DNA. This movement positions the E2 active site near the ordered lysine K120, explaining why this residue is ubiquitylated in the *C. elegans* complex (Figure S10c). Distance analysis between K120 and the E2 active site further supports this interpretation, showing a shorter distance in the DNA-anchored state relative to the DNA-uncoupled state (Figure S11a). Notably, the distance between the ordered lysine K119 and the E2 active site in human is significantly greater than the corresponding distances in *C. elegans* (Figure S11a), indicating that, in the human complex, the ordered lysine is not ubiquitylated likely due to the absence of the loop-driven movement of BCBD toward H2A C-terminal tail. In DNA-uncoupled state, the CeBCBD complex moves away from the H2A C-terminal region, with BRD-1 K54 and R55 showing reduced NCP contacts (Figure 3b,c). This repositioning elevates the E2 enzyme relative to the nucleosome (Figures S10a and S11d) and allows the H2A tail to swing upward, enabling ubiquitylation of H2A K125 within the disordered region. In this configuration, the tail has greater freedom of motion and can approach the E2 active site (Figure S10a). This state closely resembles the predominant motion in the human complex and may explain why ubiquitylation in the human system occurs primarily on disordered lysines. Close proximity below ∼ 9 Å between the disordered lysines and the E2 active site was not observed in our simulation (Figure S11b). In the DNA-proximal state the distance remains comparable to that in the DNA-uncoupled state which may be due to the combination of *Tilt-1* and *Tilt-2* motion, placing the E2 active site at similar distances from K120, but in opposite orientations relative to one another (Figure S11c,d). Figure S12a shows the similarity between the *C. elegans* E3 and E2 in DNA-uncoupled state and the human cryo-EM structure (PDB 7JZV^12^), whereas Figure S12b,c highlights the differences between human and *C. elegans* DNA-proximal and DNA-anchored states, respectively. Together, these results indicate that the BRD-1 loop provides the *C. elegans* complex with two distinct modes of motion: one that favors ubiquitylation of the ordered lysine (K120) and another that targets the disordered lysine (K125). In human, the lack of the former motion restricts ubiquitylation to the ordered region, leading to more selective modification of the disordered lysine only.

## Discussion

Our comparative analysis of the HsBCBD and CeBCBD complexes shows that an apparently small architectural difference, specifically an extra loop insertion in BRD-1, alters nucleosome engagement and influences ubiquitylation specificity. Despite conserved RING domains and intact E3 activity in both complexes, this DNA-interacting loop expands accessible motions of the CeBCBD complex and alters how the E2 active site is positioned relative to H2A lysines.

The additional loop in BRD-1 is the principal structural difference from human BARD1 and accounts for the principal divergence within the BCBD. This is not unlike comparisons between BARD1 and the non-E2 binding partner of the PRC1 E3 ligase, BMI1, which also contains an additional loop that forms contacts with the nucleosome. ^55^ However, in the case of BMI1, the loop is located in a different region of the RING domain and basic residues on the loop interact with acidic residues on H3 and H4. In the *C. elegans* complex, this loop promotes a contact pattern with NCP that is broader in extent but weaker per interaction, consistent with a more dynamic, transient engagement with the nucleosome. Like PRC1, the dominant differences map to the BRD-1/BARD1/BMI1 partner rather than main E2-binding domain of the heterodimer (BRCA1/BRC-1/RING1b), implicating the accessory subunit as the main source of variation. Within the BRD-1 loop, residues K54 and R55 form stable DNA contacts, identifying this region as a key nucleosome-binding element. Together, these features suggest loop insertions may drive differences in homolog and ortholog binding modes that are likely to influence downstream ubiquitylation.

Dynamic analyses and mutagenesis support this interpretation. Mobility is large in the BRD-1 loop, and the *C. elegans* complex adopts more tilted orientations than the human complex. Principal component analysis indicates that, in addition to a shared dominant motion, BRD-1 enables two further, more pronounced modes along PC2 that are uncommon in the human complex. These worm-specific modes correlate with K54/R55 contacts to the nucleosome, and electrostatics and vdW interaction energies indicate that the interaction is primarily DNA-mediated rather than histone-mediated. Likewise, mutagenesis of K54 and R55 to negatively charged amino acids results in a significant loss of nucleosome-ubiquitylation activity.^14^ Taken together, the results are consistent with a loop-mediated “engage-and-tilt” mechanism in which DNA engagement by K54/R55 promotes tilting and broadens the conformational ensemble.

One factor contributing to the increase in dynamics in the worm system could be evolution at an overall lower temperature than the human complex. Comparison of protein dynamics of enzymes evolved at different temperatures using either solution experiments or molecular dynamics established that enzymes from thermophiles have less dynamics and more stability compared to their mesophile homologs. ^56–59^ Our results suggest that a similar trend, where protein dynamics are inversely related to the optimal temperature of the organism, may hold even within mesophile homologs; *C. elegans* thrives in soil temperatures around 20 °C, while humans maintain a higher body temperature of 37 °C. The stable BCBD-NCP complex dictates that the human system relies on rapid H2A tail motions for lysine encounters with the active site. We did not observe a close encounter (*<* 8.98 Å) for human in our three simulations for 2400 ns at 30°C (Figure S11b), indicating these motions could be rate-limiting. Increased temperature will provide for more rapid H2A tail motions making ubiquitylation possible, while the *C. elegans* complex may require the increased motion of the E2 active site at lower temperatures to balance the slower-moving H2A tail. In line with this interpretation, Figure S11b shows that the disordered H2A tail lysine in the *C. elegans* DNA-uncoupled state reaches similarly short distances to the E2 active site as in the human system, while remaining in this proximity for a longer time, potentially because 30°C is higher than the worm’s typical soil temperature but lower than normal human body temperature. Additional specificity and stability could also be provided to the *C. elegans* complex via domains of BRC-1 and BRD-1 outside of the RING domains. Unfortunately, these domains are not present in our *in vitro* or *in silico* experiments because including them would be cost and time-prohibitive.

Changes in geometry of the *C. elegans* complex also directly impacts ubiquitylation. In *C. elegans* DNA-proximal and DNA-anchored states, where K54/R55–DNA contacts are robust, CeBCBD bends toward the H2A C-terminal region, and particularly in DNA-anchored state, positions the E2 active site near the ordered lysine K120, favoring its ubiquitylation. In DNA-uncoupled state, shared by both species, the complex shifts toward the N-terminal region; contacts between the BRD-1 extra loop and NCP diminish, the E2 is elevated, and the H2A tail gains freedom to move upward, favoring modification of a disordered lysine (K125). Because HsBCBD predominantly occupies this state, our data explains why the human complex mainly modifies disordered lysines, whereas CeBCBD alternates between two productive modes: one targeting ordered K120 and another targeting disordered K125. It is unknown whether the differences in H2A lysine specificity affect downstream regulation of gene transcription. Human PRC1 and BCBD act on distinct C-terminal tail lysine residues, yet the activity of both complexes is associated with gene repression. ^13,60–62^ Decreases in expression have also been associated with *C. elegans* PRC1 and BCBD even though their lysine targets may be less distinct.^14,63,64^ Although recently, it was shown that loss of PRC1 function in *C. elegans* leads to both increases and decreases in expression of genes enriched with H2A ubiquitylated on K119, suggesting H2A-Ub may act on transcription through multiple mechanisms depending on the chromatin structure and surrounding modifications.^65^ Typically, readers of modified histones recognize not just the modification, but residues on the nucleosome surrounding the modification to provide specific downstream signals unique to each epigenetic mark. Current understanding indicates this is also the case for C-terminal tail ubiquitylation: In humans, SMARCAD1 has been proposed as a reader for H2A-Ub on the far C-terminal Lys and ZRF1 the reader for C-terminal lysines closer to the H2A core (reviewed in^66^). However, not enough structural information is available to determine how or if ZRF1 and SMARCAD1 recognize H2A-Ub sequence specifically. Both readers have *C. elegans* homologs, providing two divergent but semi-conserved systems to further investigate mechanisms of gene repression driven by H2A-Ub.

To explore whether the relationship between BARD1/BRD-1 loop length and H2A tail length reflects a broader evolutionary trend beyond the two model organisms, we extended the analysis across Nematoda and Mammalia. The reciprocal relationship holds throughout, in which all *Caenorhabditis* species display a longer BRD-1 loop insertion paired with a shorter H2A tail, while all mammalian species show the opposite, a longer H2A tail and no BARD1 insertion (Figure S13). However, among the deposited BARD1 homolog sequences within these groups, only *Caenorhabditis* (worm) species contained a substantial loop insertion, with no intermediate cases identified elsewhere. This lack of variation in loop length limits our ability to fully evaluate a compensatory elongation pattern with confidence.

By resolving the structural mechanisms that bias Ub transfer in each complex, our study highlights the value of comparing complexes from different species with conserved functions.

Our findings raise the possibility that BCBD participates in chromatin-regulatory pathways, and that its contribution may vary depending on genomic regions, cell types, or species, even though its overall function is conserved. Future work should test how BCBD-driven lysine choice is influenced by the chromatin environment, histone variants, and whether similar nucleosome-engagement principles apply to other RING E3 ligases.

## Data and Software Availability

The input parameter and coordinate files for MD simulations are available at Zenodo (DOI: https://doi.org/10.5281/zenodo.19474007).

## Author Contributions

S.H.: conceptualization, *in silico* investigation, writing and editing, data visualization; C.L.: protein preparation, mutagenesis, and H2A ubiquitylation assays; L.H.: protein preparation and H2A ubiquitylation for mass spectrometry; T.S.: *in silico* investigation, data visualization; F.H.: Coevolutionary analysis, F.M.: supervision of Coevolutionary analysis; S.T.W.: mass spectrometry data collection, analysis, and presentation; H.T. and M.D.S: conceptualization, data visualization, writing and editing, supervision, funding acquisition, project administration.

## Acknowledgement

The authors thank Rachel Klevit, Ph.D. and Sam Witus, Ph.D. at University of Washington for purifying octomers containing histone tail mutations used in this work. This work was supported by National Institutes of Health grants R35 GM155106 (H.T.), R15 GM135900 (M.D.S.), R35 GM133631 (F.M.), and P30 CA54174 (UTHSCSA Institutional Mass Spectrometry Laboratory), as well as a startup grant from the University of Texas at Dallas to H.T. We acknowledge the Texas Advanced Computing Center (TACC) at the University of Texas at Austin and the Office of Information Technology and Cyber Infrastructure Research Computing (CIRC) department and the High Performance Computing at the University of Texas at Dallas (HPC@UTD) for their technical assistance and providing the computing resources that have contributed to this research. Mass spectrometry analyses were conducted at the University of Texas Health Science Center at San Antonio (UTHSCSA) Institutional Mass Spectrometry Laboratory, with expert technical assistance of Sammy Pardo and Dana Molleur, under the direction of Susan T. Weintraub, supported in part by UTHSCSA and by the University of Texas System Proteomics Core Network for the purchase of the Orbitrap Fusion Lumos mass spectrometer. Publication charges for this article were supported by the TCU Library Open Access Fund.

