## Supplementary material for "Small Structural Variations, Large Functional Consequences: Comparative Analysis Reveals Structural Control of Ubiquitylation Site Selection by BRCA1/BARD1": SI

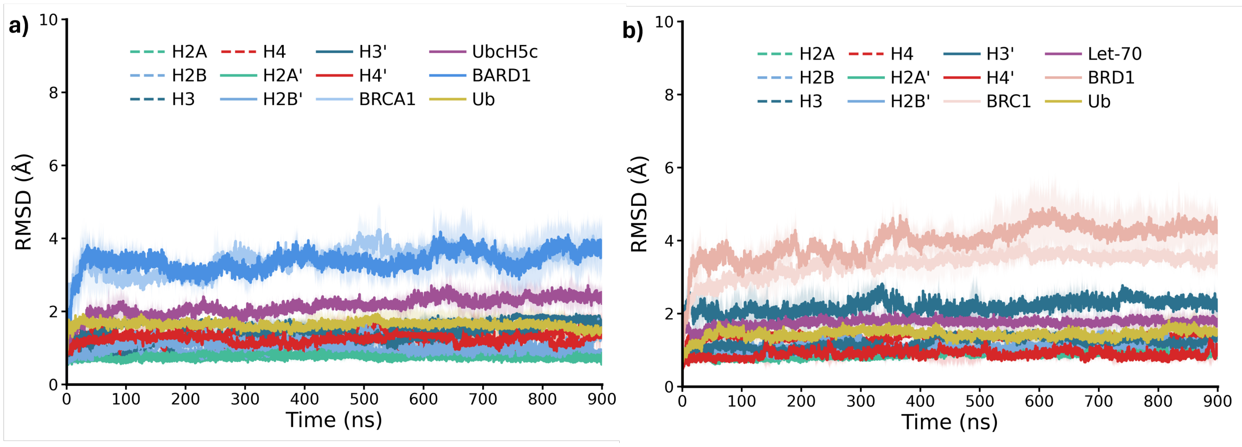


Figure S1: Average RMSD of E3, E2, Ub, and histones without tails for a) human and b) *C.elegans* systems over all simulation trials. All shading indicates the standard deviation of the data set from the average.

```
Worm    1   SGRGK-GGKAKTGGKAKSRSSRAGLQFPVGRLHRILRKGNYAQRVGAGAPVYLAAVLEY
Human   1   SGRGKQGGKA--RAKAKSRSSRAGLQFPVGRVRRLLRKGNYAERVGAGAPVYLAAVLEY

Worm    59   LAAEVLELAGNAARDNKKTRIAPRHLQLAVRNDEELNKLLAGVTIAQGGVLPNIQAVLLP
Human   58   LTAEILELAGNAARDNKKTRIIPRHLQLAIRNDEELNKLLGKVTIAQGGVLPNIQAVLLP

Worm    119   KKTGGDKE----
Human   118   KKTESHHKAKGK
```

Figure S2: Alignment of H2A sequences from human and *C. elegans* with the folded core highlighted in gray, Lys residues mutated in this study highlighted in blue, and Lys residues ubiquitylated by human BRCA1 highlighted in green.

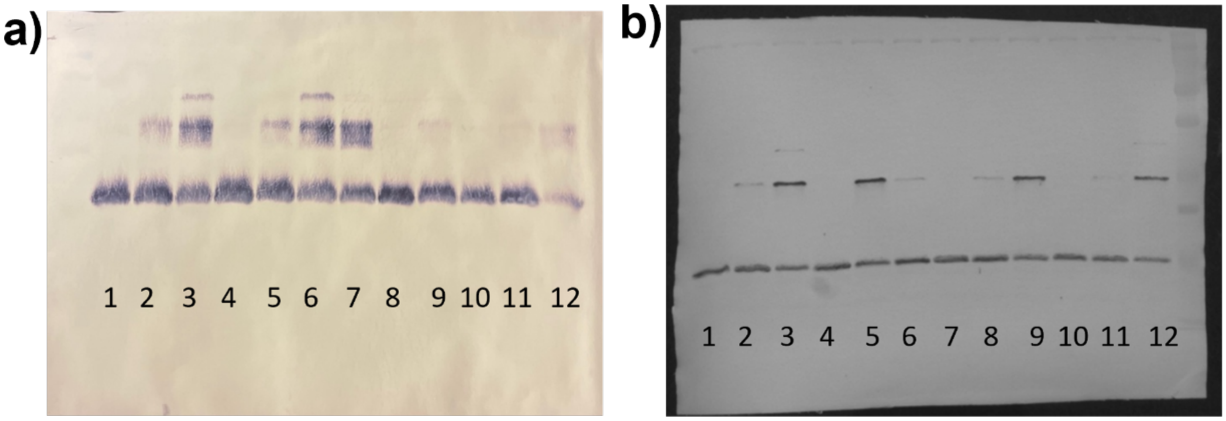


Figure S3: Uncropped, unedited blots used for quantification of H2A ubiquitylation reported in main text figure 1. H2A ubiquitylation with *C. elegans* BRCA1 was performed in duplicate using various constructs of H2A. All mutations are made relative to the H2A hybrid using the native *C. elegans* C-terminal sequence on the human H2A core and N-terminus. a) Lanes contain the following: H2A hybrid assayed for 0 min (1), 10 min (2), or 30 min (3); H2A with N-terminal Lys 14 and 16 mutated to Arg assayed for 0 min (4), 10 min (5), or 30 min (6); H2A with C-terminal Lys 119 and 120 mutated to Arg and Lys 125 mutated to His assayed for 30 min (7), 0 min (8), or 10 min (9); and all five mutation sites combined assayed for 0 min (10), 10 min (11), and 30 min (12). b) Lanes contain the following: H2A hybrid assayed for 0 min (1), 10 min (2), or 30 min (3); H2A with all C-terminal Lys mutated as described above assayed for 0 min (4), 30 min (5), or 10 min (6); all five mutation sites combined assayed for 0 min (7), 10 min (8), and 30 min (9); and H2A with N-terminal Lys mutated assayed for 0 min (10), 10 min (11), or 30 min (12).

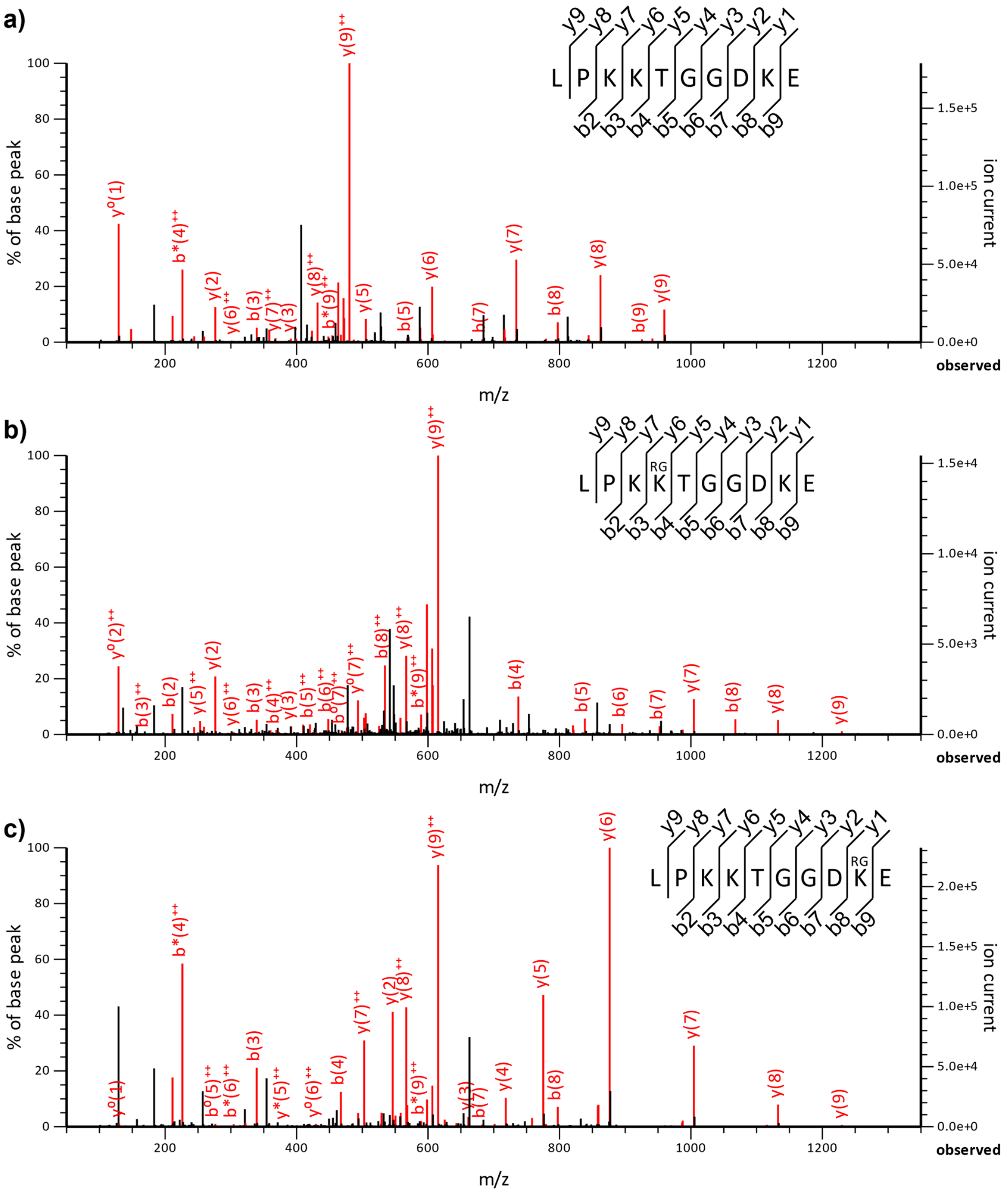


Figure S4: Tandem mass spectra (MS/MS) of ubiquitylated H2A after digestion with chymotrypsin. a) The unmodified C-terminal H2A tail is shown for comparison with two observed modified peptides (b and c) which document the residual RGG of the C-terminal ubiquitin fragment after chymotrypsin cleavage after L in the LRGG C-terminus. (RGG is shown here as RG due to space constraints.) The expected correspondence in mass of b and y ions can be seen by comparing the spectra in b and c with the unmodified peptide in a.

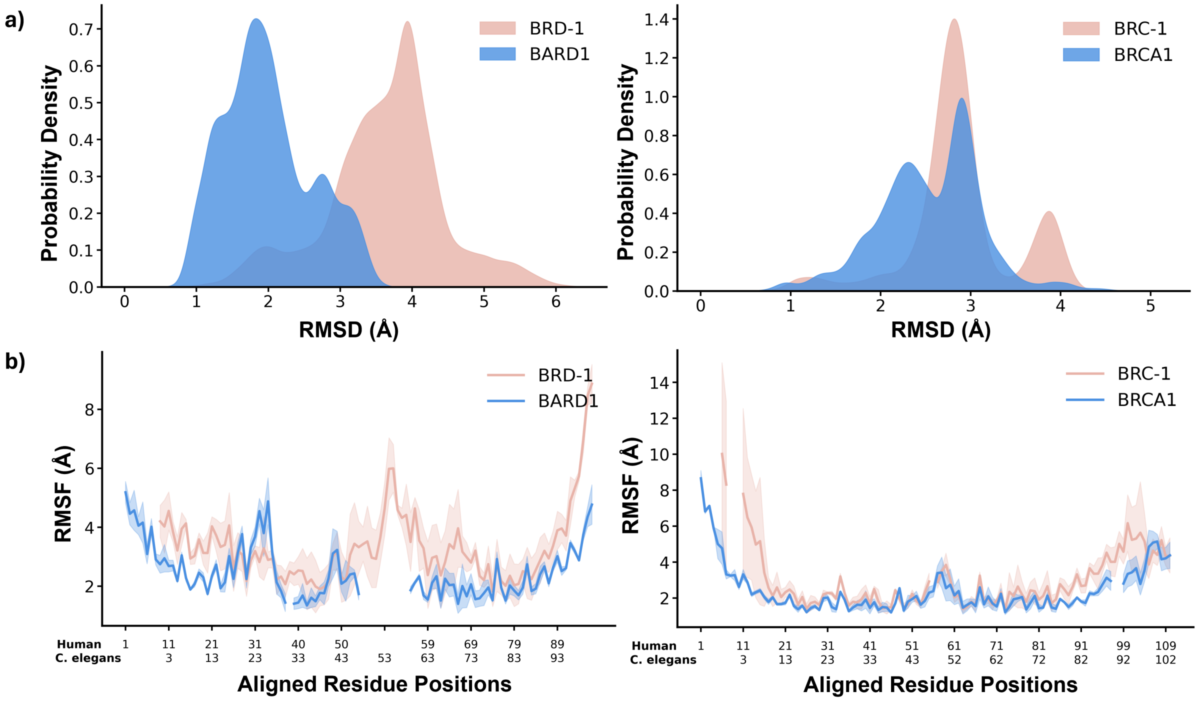


Figure S5: a) Comparison of RMSD distributions between BRD-1/BARD1 and BRC-1/BRCA1 over all trials; b) Average RMSF profiles for BRD-1/BARD1 and BRC-1/BRCA1. The x-axis shows residue numbers aligned between human and *C. elegans*.

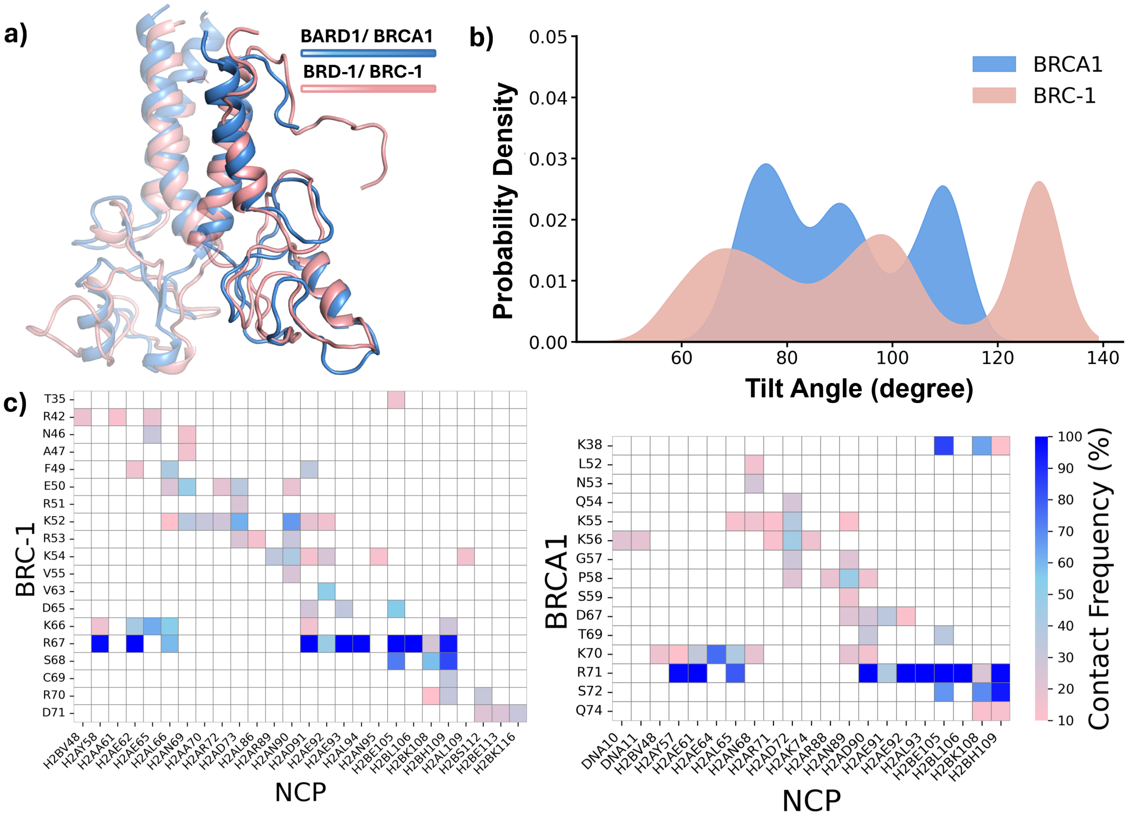


Figure S6: a) Comparisons of minimized structures (nucleosome-bound HsBCBD (PDB ID: 7JZV) vs minimized structure of nucleosome-bound CeBCBD). b) Tilt-angle analysis for BRCA1 and BRC-1. b) Differences in BRCA1 vs BRC-1 nucleosome contacts over the course of simulation analyzed by contact frequency.

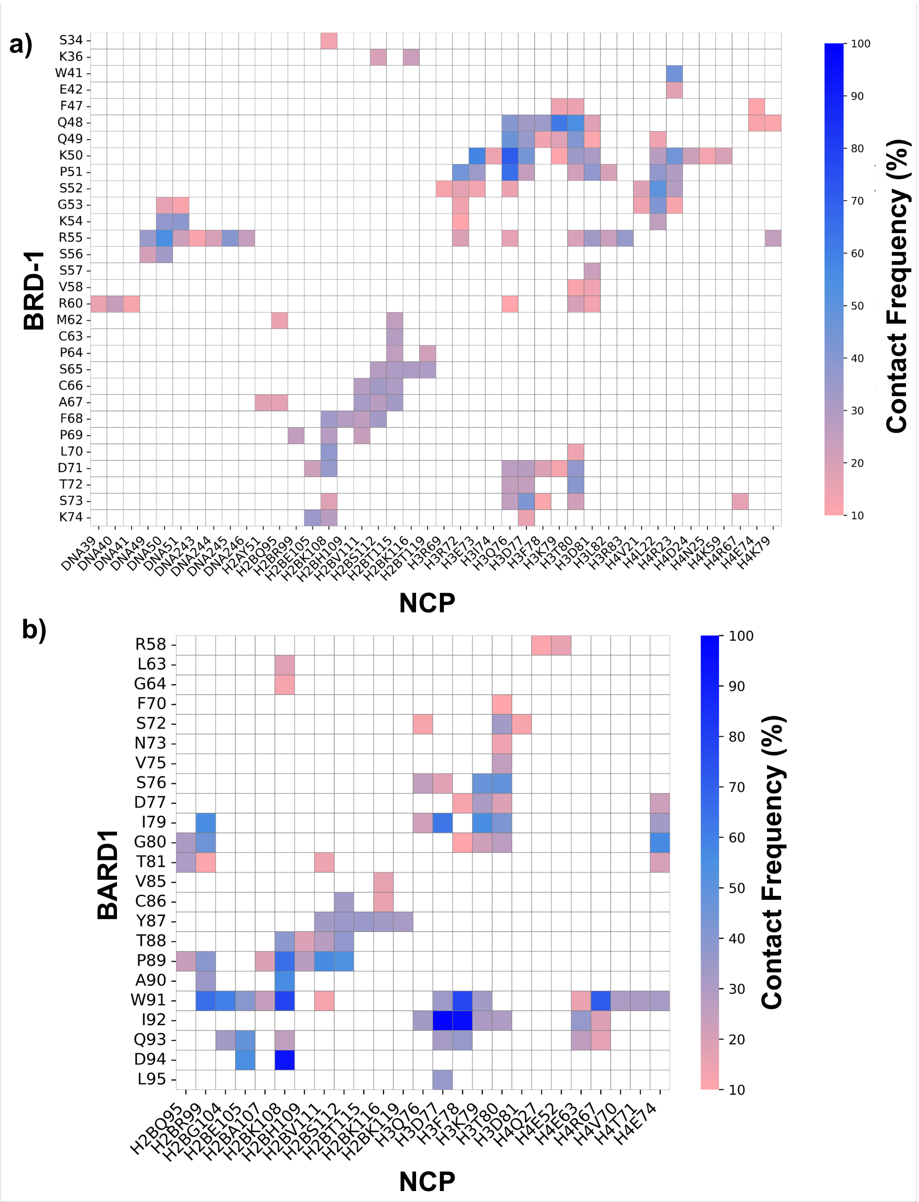


Figure S7: a) BRD-1 and b) BARD1 nucleosome contacts over the course of simulation, analyzed by contact frequency.

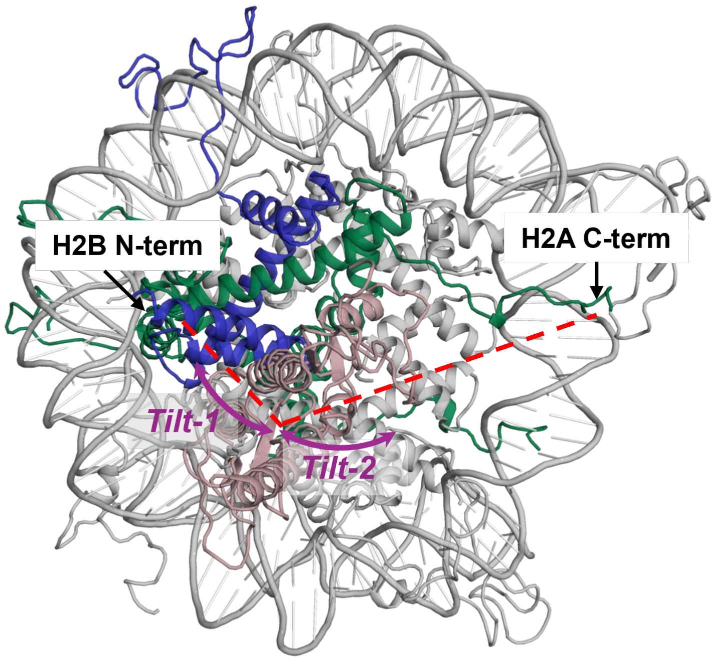


Figure S8: Top-view schematic representation of the NCP and the *C. elegans* E3 complex, showing *Tilt-1* and *Tilt-2*, corresponding to PC1 and PC2, respectively. H2A, H2B, and the *C. elegans* E3 complex are colored green, blue, and light pink, respectively. Red dashed lines indicate the orientation of the E3 complex relative to the H2B N-terminal and the H2A C-terminal tail.

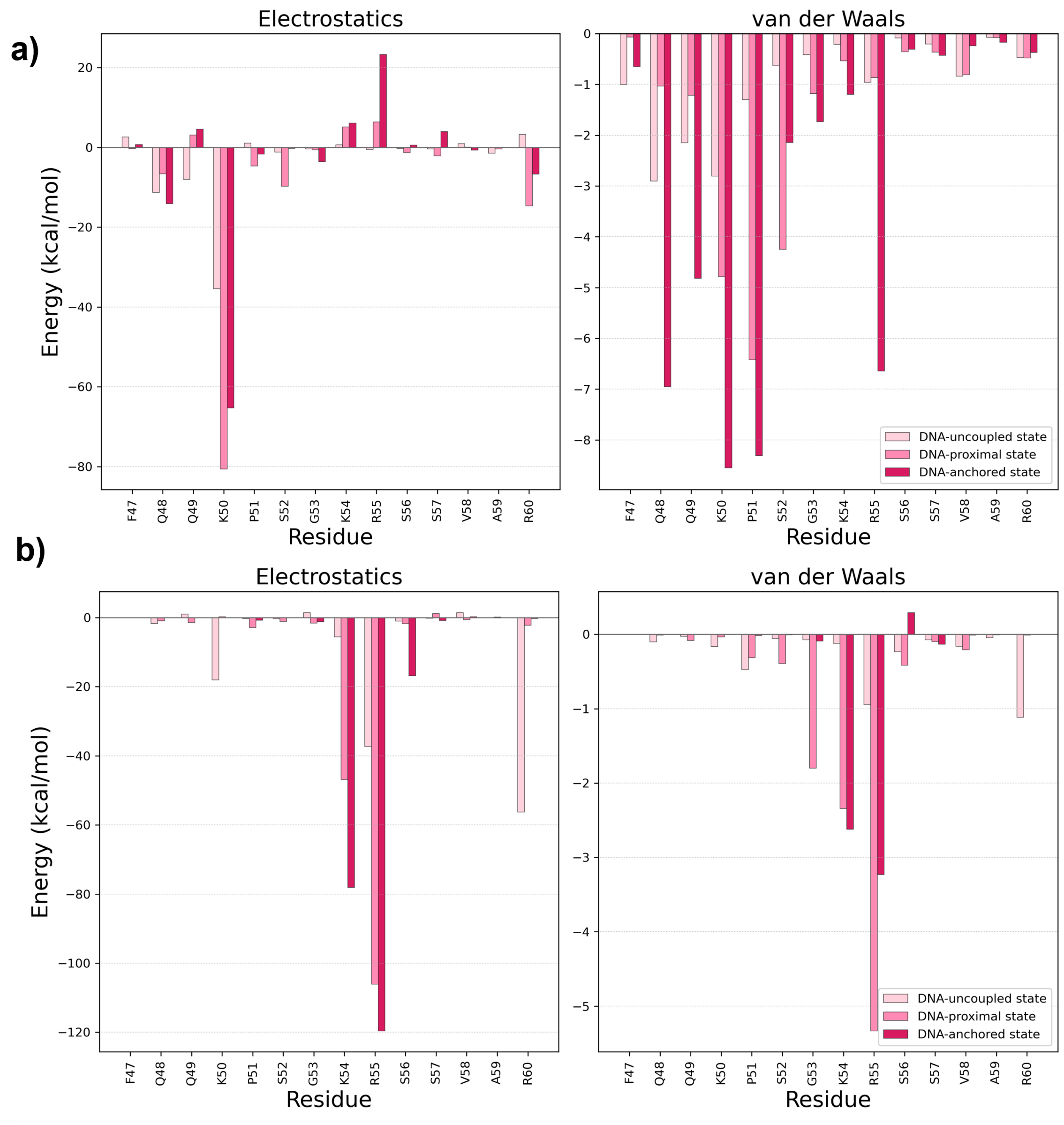


Figure S9: Electrostatic and van der Waals interaction energies between the BRD-1 extra loop and (a) histones and (b) DNA, computed via the linear interaction energy (lie) analysis.

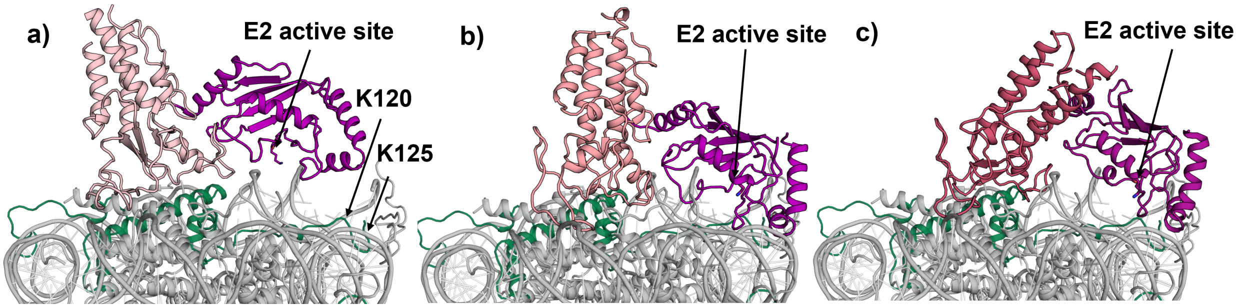


Figure S10: *C. elegans* conformational states: a) DNA-uncoupled state, b) DNA-proximal state, c) DNA-anchored state. The E3 is shown in distinct shades of pink, the E2 in purple, H2A in green, and the remaining NCP in silver.

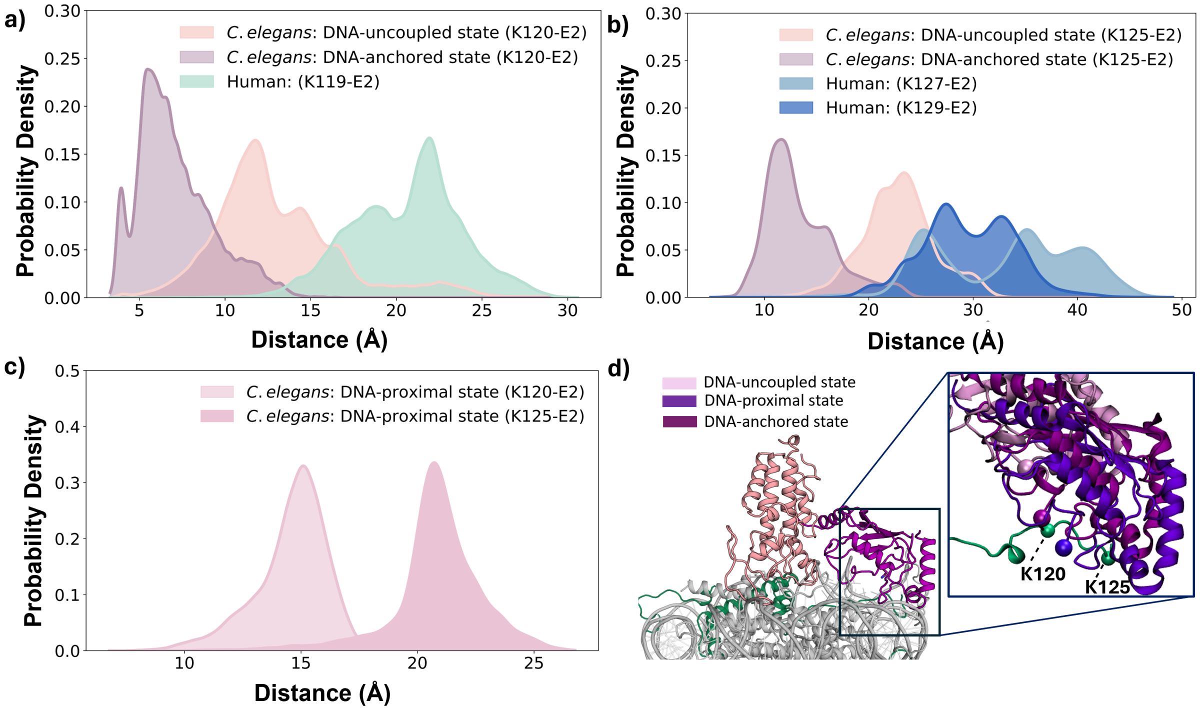


Figure S11: Distances between lysine residues targeted for ubiquitylation and the E2 active site in human and *C. elegans* across different conformational states. Comparison of distances from a) ordered and b) disordered target lysine residue(s) to the E2 active site in human and in the DNA-uncoupled and DNA-anchored states of *C. elegans*. c) Distances from ordered and disordered target lysine residue(s) to the E2 active site in the DNA-proximal state of *C. elegans*. d) Schematic representation of H2A-tail lysine residues and the E2 active site from one snapshot of aligned structures across different states, shown in different shades of purple.

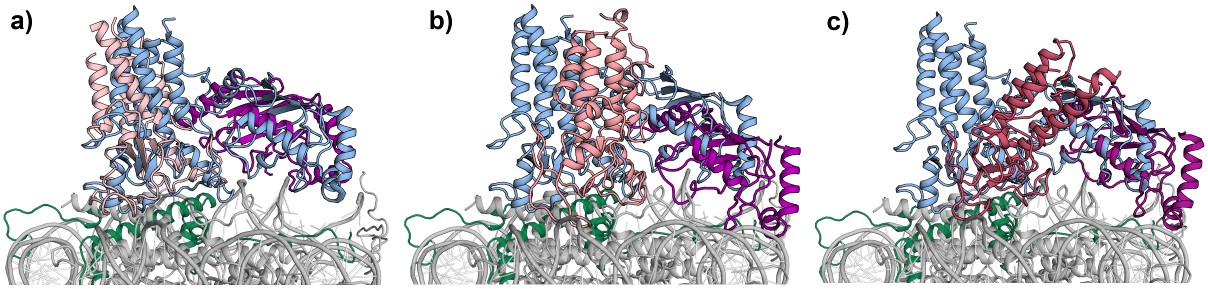


Figure S12: Superimposed structures of the *C. elegans* E3-E2 complex in a) DNA-uncoupled state, b) DNA-proximal state, and c) DNA-anchored state and human cryo-EM structure. E3 and E2 in *C. elegans* are in pink and purple shades, respectively, while E3-E2 in human is in blue, H2A in green, and the remaining NCP in silver.

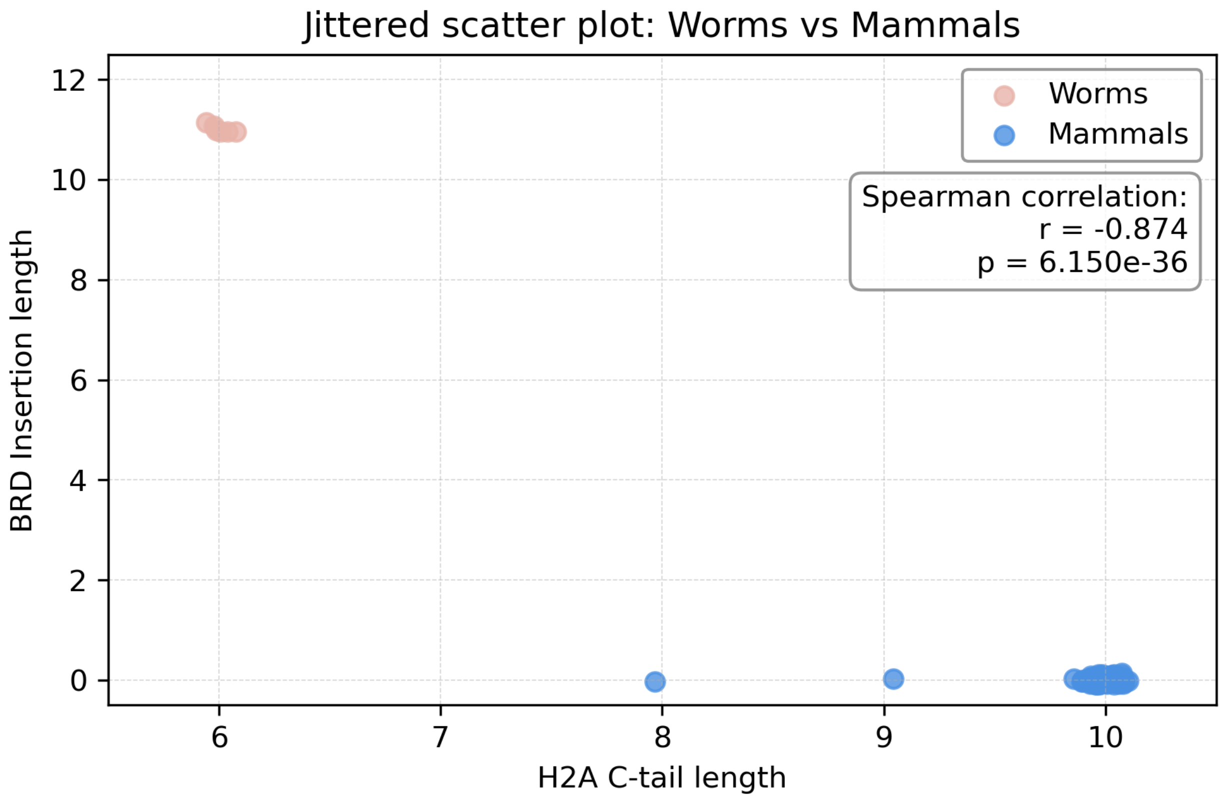


Figure S13: The jittered scatter plot depicts the association between BARD1/BRD-1 insertion length and H2A C-terminal tail length across a broad sample of species in the phylum Nematoda (worms) and the class Mammalia (mammals). In worms, presence of BRD1 insertion correlates with shorter H2A tails, whereas in mammals the pattern is reversed.
